## Supplementary material for "Termination of non-coding transcription in yeast relies on both a CTD-interaction domain and a CTD-mimic in Sen1"

**List of supplementary material:**

-Figures S1-6.

-Tables S1, S2, S11, S12, S13 and S14.

-Supplementary references.

**Material provided as separate files:**

-Table S3: quantification of the readthrough index and termination defects at CUTs in the strains expressing different versions of Sen1.

-Table S4: quantification of the readthrough index and termination defects at snoRNAs in the strains expressing different versions of Sen1.

-Table S5: complete annotation of snoRNAs transcription termination sites used in this study.

-Table S6: complete annotation of CUTs transcription termination sites used in this study.

-Table S7: subset of mRNAs with quartiles employed in analyses of RNAPII distribution using the mean as the summarizing function.

-Table S8: subset of CUTs with quartiles employed in analyses of RNAPII distribution using the mean as the summarizing function.

-Table S9: subset of mRNAs with quartiles employed in analyses of Sen1 distribution using the mean as the summarizing function.

-Table S10: subset of CUTs with quartiles employed in analyses of Sen1 distribution using the mean as the summarizing function.

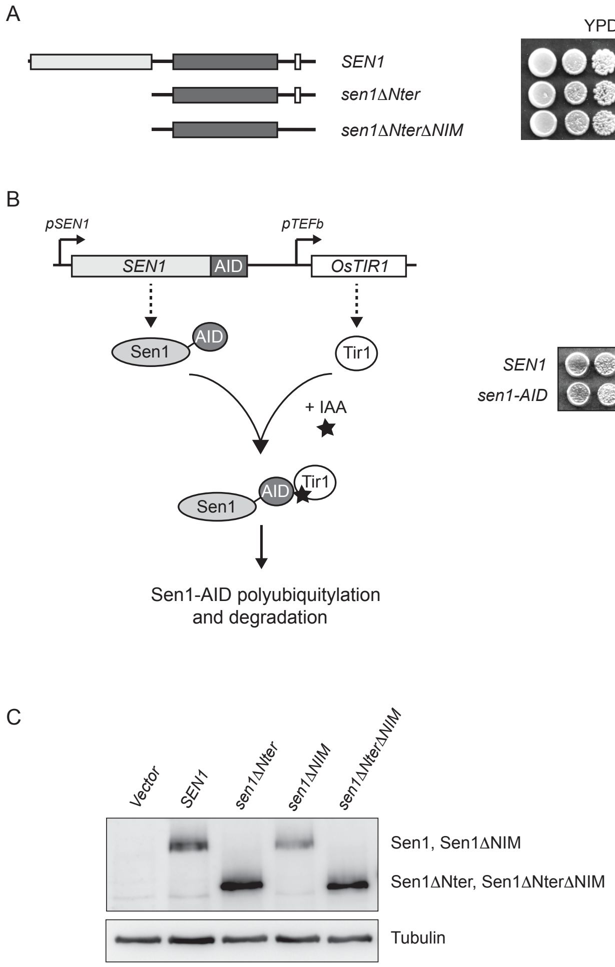


**Figure S1 (related to figure 3): A**) Growth tests performed as in figure 3A but using strains harbouring the TAP-tagged versions of *SEN1* indicated on the left at the endogenous locus in the presence of a *URA3* plasmid (pFL38) expressing *SEN1*. **B**) Description of the degron system employed in this study to deplete Sen1. The *SEN1* endogenous locus is modified by inserting the sequence of an AID (auxin inducible degron) tag followed by a cassette for the expression of *Oryza sativa TIR1* (*OsTIR1*), an ubiquitin ligase. In the presence of auxin hormones such as indole-3-acetic acid (IAA), Tir1 recognizes the AID-tag and promotes polyubiquitylation and subsequent degradation of Sen1-AID by the proteasome. A growth test illustrating the inhibition of cell growth upon depletion of Sen1 is shown on the right. **C**) Western blot analysis of HA-tagged versions of Sen1 expressed from a centromeric plasmid in a Sen1-AID strain upon depletion of the chromosomally-encoded copy of Sen1 as in figure 3B. Tubulin is detected as a loading control.

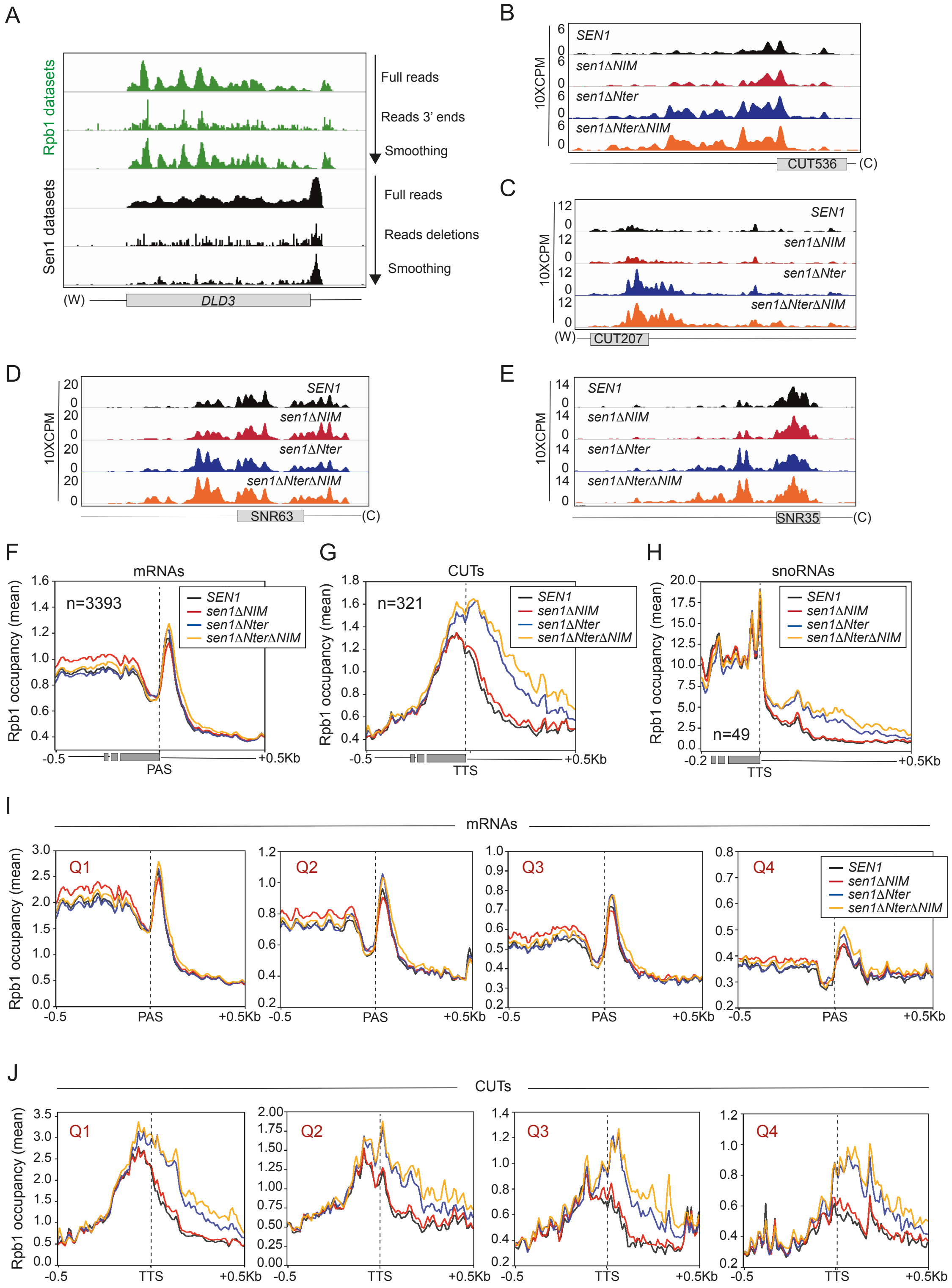


**Figure S2 (related to figure 3): A)** Screenshot of a particular mRNA to illustrate the conversions operated on RNAPII (Rpb1) and Sen1 CRAC datasets to increase resolution. **B**) and **C**) Additional examples of CUTs exhibiting transcription termination defects upon deletion of the Nter that are exacerbated in the double *ΔNterΔNIM* mutant. **D**) and **E**) Additional examples of snoRNAs showing transcription termination defects in Sen1 mutants. **F-H)**

Metagene analysis of the RNAPII distribution at the indicated classes of RNA in the presence of different Sen1 variants. Values on the y axis correspond to the mean coverage. Similar metagene analyses performed on quartiles for mRNAs (**I**) and CUTs (**J**), where Q1 represents the 25% of RNAs of the indicated class exhibiting the highest RNAPII occupancy in the wt.

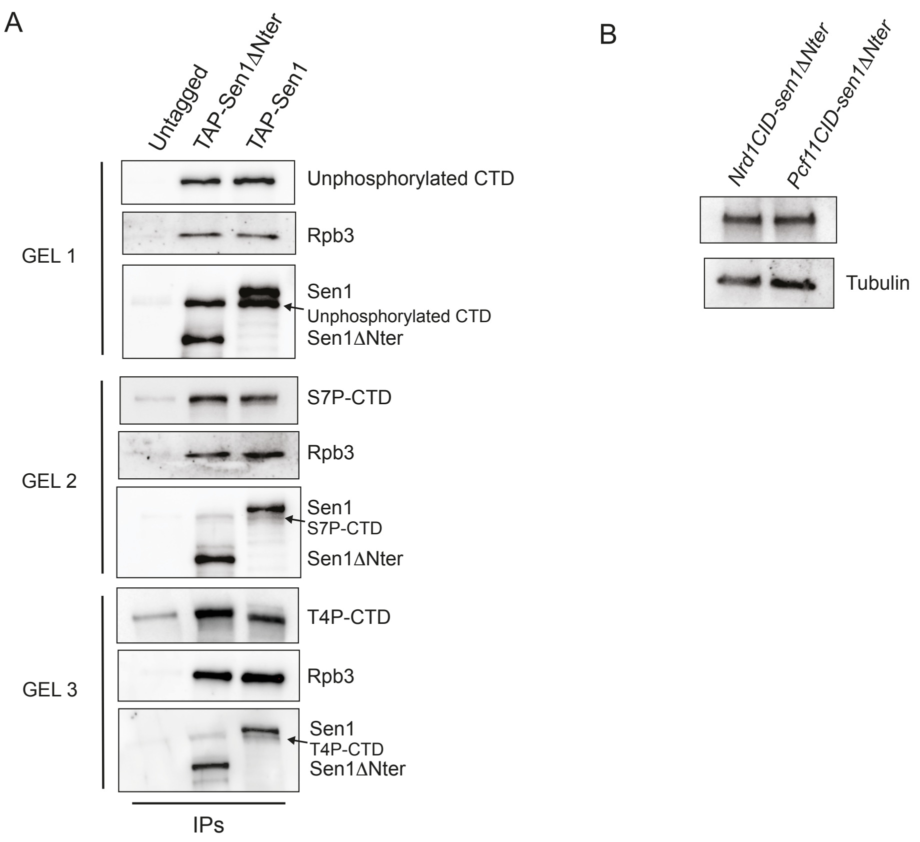


**Figure S3 (related to figure 5): A)** Deletion of the Sen1 N-terminal domain does not affect the interaction of Sen1 with the S7P-CTD or with the unphosphorylated CTD. CoIP experiments using TAP-Sen1 as the bait. Sen1 proteins were expressed from pGAL in the presence of galactose. The Rpb3 subunit of RNAPII is detected as a control. Protein extracts were treated with RNaseA before immunoprecipitation. Samples correspond to the same experiment and were loaded in three different gels run. **B)** Western blot analysis of HA-tagged versions of the chimeric Sen1 proteins indicated from a centromeric plasmid in a Sen1-AID strain upon depletion of the chromosomally-encoded copy of Sen1 as in figure 3B. Tubulin is detected as a loading control.

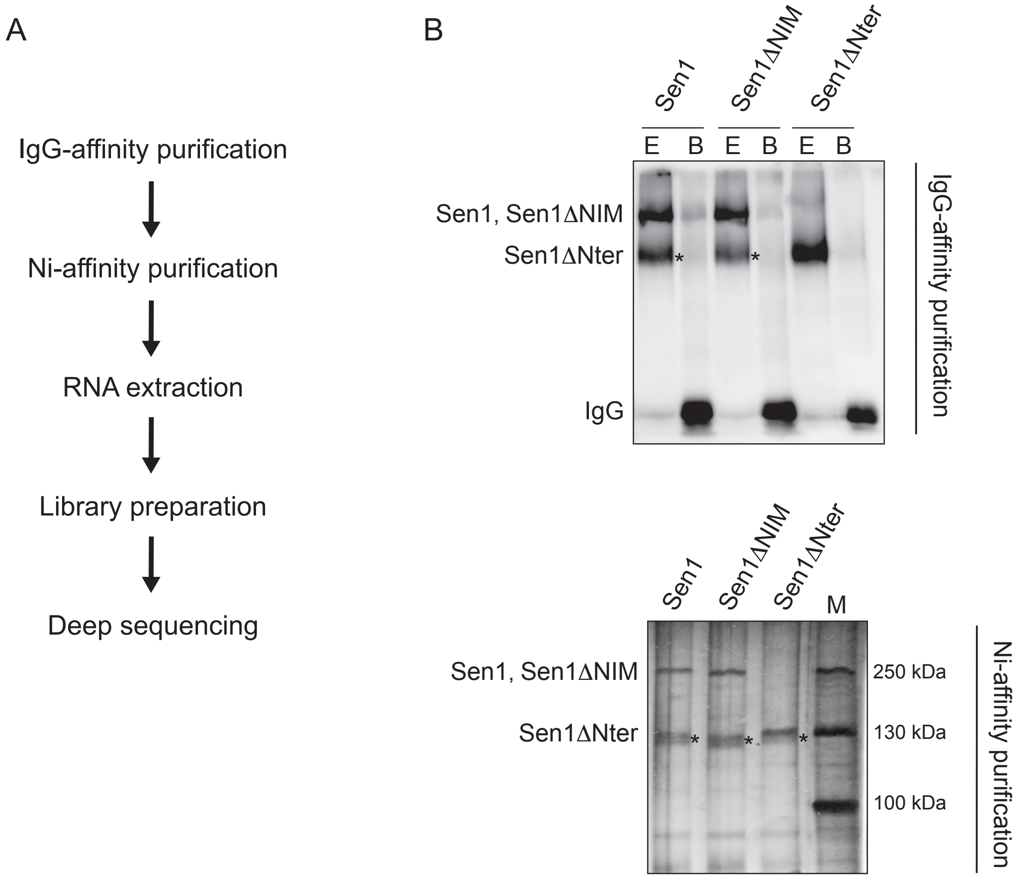


**Figure S4 (related to figure 6): A) Schematic of the main steps in a CRAC experiment. B)** Results of the two purification steps of the different HTP ($His_6$-TEV-ProteinA)- tagged Sen1 variants in a typical CRAC experiment. Representative gels corresponding to one out of two independent biological replicates. Top: western blot showing results of the IgG-affinity purification step. E, eluates after cleavage of the protein A moiety using the TEV protease. B, protein remaining associated with IgG beads. Proteins were revealed using the PAP antibody. Bottom: silver-stained denaturing gel showing the results of the Ni-affinity purification step. M, molecular-weight marker. An asterisk indicates Sen1 proteolytic fragments of size similar to that of Sen1ΔNter that we typically obtain in all Sen1 purifications.

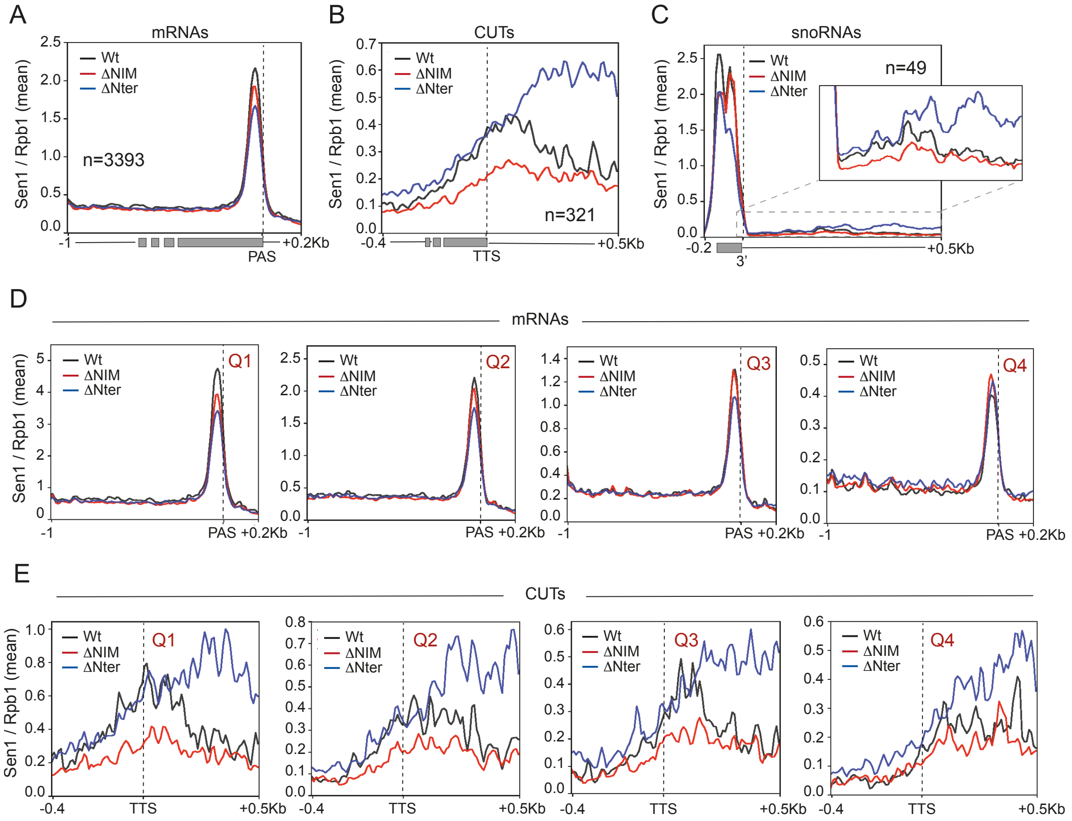


**Figure S5 (related to figure 6): A-C)** Metagene analysis of the normalized distribution of different Sen1 variants at the indicated classes of RNA. Values on the y axis correspond to the mean coverage. Similar metagene analyses performed on quartiles for mRNAs (**D**) and CUTs (**E**), where Q1 represents the 25% of RNAs of the indicated class exhibiting the highest wt Sen1 occupancy.

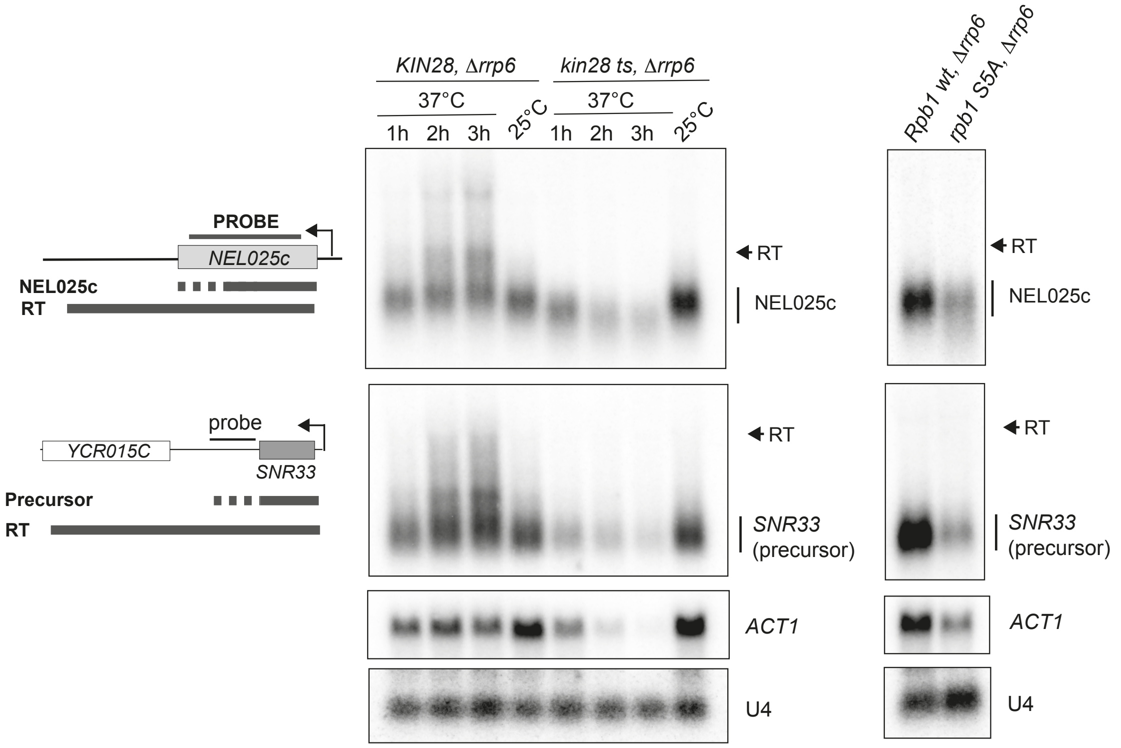


**Figure S6: Analysis of the role of S5 phosphorylation for NNS-dependent transcription termination.** Northern blot analyses of two well-characterized NNS-targets in either a *kin28* thermosensitive (ts) mutant (left) or a Rpb1-anchor away strain carrying a plasmid expressing either the wt or S5A versions of *RPB1* (right). Kin28 was inactivated by incubation at 37°C for the indicated times. Note that growth at 37°C induces mild termination defects in the control strain due to the presence of the *Δrrp6* mutation, which also confers a thermosensitive phenotype. The Rpb1-anchor away strains were treated for 2h with rapamycin to induce the nuclear depletion of the chromosomally-encoded Rpb1. The intermediate phenotype of this strain compared to the *kin28 ts* suggests incomplete depletion of the wt version of Rpb1 during the timeframe of the experiment. The expected RNA species resulting from inefficient termination (RT for readthrough species) are indicated. The *ACT1* mRNA is detected to provide a readout of the general decrease in transcription provoked by the inhibition of S5 phosphorylation. U4 RNA is used as a loading control. Representative gels of one out of two independent experiments. Probes used for RNA detection are described in table S14.

**Table S1:** Equilibrium binding of Nrd1 CID to Sen1 and Trf4 NIMs monitored by fluorescence anisotropy.

| | Sen1 NIM | | | Trf4 NIM | | |
|---|---|---|---|---|---|---|
| | Kd (μM) | | | Kd (μM) | | |
| **wt** | 1.2 | ± | 0.02 | 0.9 | ± | 0.02 |
| **L20D** | 16.9 | ± | 1.5 | 13.4 | ± | 1.2 |
| **K21D** | 8.0 | ± | 0.8 | 6.1 | ± | 0.4 |
| **S25D** | 76.1 | ± | 14.2 | 53.9 | ± | 8.2 |
| **R28D** | 78.2 | ± | 11.5 | 63.4 | ± | 4.0 |
| **I130A** | 7.6 | ± | 0.3 | 4.5 | ± | 0.2 |
| **I130K** | 36.7 | ± | 4.6 | 14.1 | ± | 1.2 |
| **I130N** | 8.9 | ± | 0.5 | 6.1 | ± | 0.3 |
| **M126A** | 1.4 | ± | 0.07 | 1.1 | ± | 0.02 |
| **R133A** | 1.8 | ± | 0.08 | 1.4 | ± | 0.02 |
| **R133D** | 2.0 | ± | 0.1 | 1.2 | ± | 0.05 |
| **R133G** | 1.6 | ± | 0.04 | 1.3 | ± | 0.04 |

**Table S2: NMR and refinement statistics for the Nrd1 CID–Sen1 NIM complex**

| | Nrd1 CID-Sen1 NIM complex |
|---|---|
| **NMR distance & dihedral constraints** | |
| Distance restraints | |
| Total NOEs | 2499 |
| Intra-residue | 654 |
| Inter-residue | 1845 |
| Short | 1273 |
| Medium | 645 |
| Long | 581 |
| Hydrogen bonds | 100 |
| Intermolecular distance restraints | 65 |
| Total dihedral angle restraints[a] | 222 |
| **Structure statistics**[b] | |
| Violations (mean and s.d.) | |
| Number of distance restraint violations > 0.5 Å | 0.05 ± 0.22 |
| Number of dihedral angle restraint violations > 15° | 1.1 ± 1.2 |
| Max. dihedral angle restraint violation (°) | 29.03 ± 27.63 |
| Max. distance constraint violation (Å) | 0.027 ± 0.12 |
| Deviations from idealized geometry[b] | |
| Bond lengths (Å) | 0.0037 ± 0.00008 |
| Bond angles (°) | 1.673 ± 0.013 |
| Average pairwise r.m.s.d (Å)[b] | |
| Nrd1 CID (6-82,91-135,143-147,149-153) | |
| Heavy atoms | 1.1 |
| Backbone atoms | 0.7 |
| Sen1 NIM (2052-2063) | |
| Heavy atoms | 1.9 |
| Backbone atoms | 1.3 |
| Complex | |
| All complex heavy atoms | 2.5 |
| All complex backbone atoms | 2.0 |
| Ramachandran plot statistics[c] | |
| Residues in most favoured regions (%) | 96.7 |
| Residues in allowed regions (%) | 3.2 |
| Residues in generously allowed regions (%) | 0.1 |
| Residues in disallowed regions (%) | 0.0 |

[a] α-helical dihedral angle restraints imposed for the backbone based on the CSI.
[b] Calculated for an ensemble of the 20 lowest energy structures.
[c] Based on Procheck analysis (Laskowski et al., 1996).

**Table S11**: Antibodies used in this work.

| Name | Reference/provider | Working dilution | Use |
|---|---|---|---|
| Anti-HA F7 | Sc-7392/ Santa Cruz | 1:1000 | Sen1-HA detection in figures 1A, 1C-D, 5A |
| Anti-HA 12CA5 | 11 666 606 001/ Sigma | 2.5 ug/reaction | Sen1 coIP in figure 1D |
| Anti-CBP, calmodulin binding protein | 07-482/ Millipore | 1:1000 | Nrd-CBP detection in figure 1A, Sen1-CBP detection in figure 5B |
| Anti-Air2 | S. Vanacova | 1:1000 | Air2 detection in figure 1A |
| Anti-Rpb1 y80 | Santa Cruz (not available any more) | 1:1000 | Rpb1 detection in figure 5A, 5B |
| Anti-phospho-CTD Ser5 clone 3E8 | 04-1572/ Millipore | 1:1000 | Detection of S5P-CTD of RNAP II in figure 5B |
| Anti-phospho-CTD Ser2 clone 3E10 | 04-1571/ Millipore | 1:1000 | Detection of S2P-CTD of RNAP II in figure 5B |
| Anti-phospho-CTD Ser7 clone 4E12 | 04-1570/ Millipore | 1:1000 | Detection of S7P-CTD of RNAP II in figure S3 |
| Anti-phospho-CTD Thr4 clone | 61362/Active motif | 1:500 | Detection of T4P-CTD of RNAP II in figure S3 |
| Anti-CTD 8WG16 | Ab817/abcam | 1:1000 | Detection of the unphosphorylated CTD preferentially. |
| PAP, peroxidase anti-peroxidase | P1291-1ML/ Sigma | 1:3000 | Used for Protein A detection on TAP tag. |
| Anti-Nrd1 | Covalab | 1:3000 | Nrd1 detection in figures 1 and 5 |
| Anti-Nab3 | Covalab | 1:3000 | Nab3 detection in figures 1 and 5 |
| Anti-His | H1029,Sigma | 1:2000 | $His_6$-GST-Sen1 Cter and $His_6$-GST in figure 5E |

**Table S12**: Yeast strains used in this work.

| Number | Name | Genotype | Source |
|---|---|---|---|
| DLY17 | W303 | *ura3-1, ade2-1, his3-11,5, trp1-1, leu2-3,112, can1-100* | (Thomas and Rothstein, 1989) |
| DLY467 | KIN28 | *kin28::URA, [pSF19-KIN28, CEN, TRP]* | (Cismowski et al., 1995) |
| DLY468 | kin28 ts16 | *kin28::URA, [pSF19-kin28-ts16 CEN, TRP]* | (Cismowski et al., 1995) |
| DLY671 | BMA | *as W303, Δtrp1* | F. Lacroute |
| DLY814 | *Δrrp6* | *as W303, rrp6::KAN* | (Porrua et al., 2012) |
| DLY1657 | *sen1-HA* | *as BMA, Sen1::HA::KAN* | (Tudek et al, 2014 |
| DLY2014 | *nrd1 ΔCID-TAP, sen1-HA* | *as BMA, nrd1ΔCID (Δ6-150)::TAP::HIS3* | Tudek et al, 2014 |
| DLY2769 | *sen1ΔNIM* | *as BMA, sen1ΔNIM (Δ2052-2063)* | This work |
| DLY2982 | *sen1ΔNIM-HA* | *as BMA, sen1ΔNIM (Δ2052-2063)::HA::KAN* | This work |
| DLY1737 | *nrd1-TAP, sen1-HA* | *as BMA, NRD1::TAP::HIS5 Sen1::HA::KAN* | (Tudek et al, 2014) |
| DLY2622 | *nrd1-TAP, sen1ΔNIM-HA* | *as BMA, NRD1::TAP::HIS5, sen1ΔNIM (Δ2052-2063)::HA::KAN* | This work |
| DLY2657 | *sen1ΔCter-HA* | *as BMA, sen1ΔCter (Δ1930-2231)::HA::KAN* | This work |
| DLY2658 | *nrd1-TAP, sen1ΔCter-HA* | *as BMA, NRD1::TAP::HIS5, sen1ΔCter (Δ1930-2231)::HA::KAN* | This work |
| DLY2695 | *sen1ΔCter-HA, Δrrp6* | *as BMA, sen1ΔCter (Δ1930-2231)::HA::KAN, rrp6::URA* | This work |
| DLY2694 | *sen1-HA, Δrrp6* | *as BMA, sen1::HA::KAN, rrp6::URA* | This work |
| DLY2770 | *sen1ΔNIM, Δrrp6* | *as BMA, sen1ΔNIM, rrp6::KAN* | This work |
| DLY2767 | *Δsen1*/pFL38-*SEN1* | *as BMA, sen1::KAN, harbouring plasmid pFL38-SEN1* | This work |
| DLY1656 | $P_{GAL1-}$ *TAP-SEN1* | *as BMA, TRP1::Pgal::TAP::SEN1* | (Porrua et al., 2012) |
| DLY2778 | $P_{GAL1-}$ *TAP-sen1ΔNIM* | *as BMA, TRP1::Pgal::TAP::sen1ΔNIM (Δ2052-2063)* | This work |
| DLY2692 | $P_{GAL1-}$ *TAP-sen1ΔNter* | *as BMA, TRP1::Pgal::TAP::sen1ΔNter (Δ1-975)* | This work |
| DLY2779 | $P_{GAL1-}$ *TAP-sen1ΔNter ΔNIM* | *as BMA, TRP1::Pgal::TAP::sen1ΔNter (Δ1-975) ΔNIM (Δ2052-2063)* | This work |
| DLY2060 | $P_{GAL1-}$ *TAP-SEN1, Δrrp6* | *as BMA, TRP1::Pgal::TAP::SEN1, rrp6::KAN* | (Porrua et al., 2012) |
| DLY2780 | $P_{GAL1-}$ *TAP-sen1ΔNIM, Δrrp6* | *as BMA, TRP1::Pgal::TAP::sen1ΔNIM (Δ2052-2063), rrp6::KAN* | This work |
| DLY2698 | $P_{GAL1-}$ *TAP-sen1ΔNter,Δrrp6* | *as BMA, TRP1::Pgal::TAP::sen1ΔNter (Δ1-975), rrp6::KAN* | This work |

| | | | |
|---|---|---|---|
| DLY2781 | *$P_{GAL1}$- TAP-sen1ΔNterΔNIM, Δrrp6* | *as BMA, TRP1::Pgal::TAP::sen1∆Nter (∆1-975) ΔNIM (Δ2052-2063), rrp6::KAN* | This work |
| DLY2782 | *sen1-AID* | *as BMA,SEN1-AID::KAN::OsTIR1* | This work |
| DLY2788 | *sen1-AID, Δrrp6* | *as BMA,SEN1-AID::KAN::OsTIR1, rrp6::URA* | This work |
| DLY2724 | *TAP-sen1/*pFL38-*SEN1* | *as BMA, TAP::SEN1, harbouring plasmid pFL38-SEN1* | This work |
| DLY2725 | *TAP- sen1ΔNter /*pFL38-*SEN1* | *TAP::sen1∆Nter (∆1-975), harbouring plasmid pFL38-SEN1* | This work |
| DLY2726 | *TAP- sen1ΔNterΔNIM /*pFL38-*SEN1* | *as BMA, TAP::sen1∆Nter (∆1-975) ΔNIM (Δ2052-2063), harbouring plasmid pFL38-SEN1* | This work |
| DLY3152 | *$P_{GAL1}$- TAP-SEN1, Δnrd1/ pRS316-NRD1* | *as BMA, TRP1::Pgal::TAP::SEN1, nrd1::KAN; harbouring plasmid pRS316-NRD1* | This work |
| DLY3187 | *$P_{GAL1}$- TAP-SEN1, Δnab3/ pFL38-Nab3* | *as BMA, TRP1::Pgal::TAP::SEN1, nab3::KAN; harbouring plasmid pFL38-NAB3* | This work |
| DLY3081 | *rpb3-FLAG* | *as BMA, RPB3::3xFLAG::natMX6* | This work |
| DLY3151 | *rpb3-FLAG, sen1-AID* | *as BMA, SEN1-AID:KAN::OsTIR1, RPB3::3xFLAG::natMX6* | This work |
| DLY3361 | *sen1-AID, rpb1-HTP* | *as BMA, Sen1::AID, KAN::OsTIR1, rpb1-HTP::URA* | This work |
| yFR1499 | Rpb1 anchor away + Rpb1-CTD wt-3xFLAG | *ade2-1, trp1-1, can1-100, leu2-3,112, his3-11,15, ura3, GAL, psi+, tor1-1, fpr1Δ::NAT, RPL13A-2xFKBP12::TRP1, RPB1-FRB::KanMX6, [pFR482 (CEN, HIS3, RPB1-CTDWT-3xFLAG)]* | (Collin et al., 2019) |
| yFR1503 | Rpb1 anchor away + Rpb1-CTD S5A-3xFLAG | *ade2-1, trp1-1, can1-100, leu2-3,112, his3-11,15, ura3, GAL, psi+, tor1-1, fpr1Δ::NAT, RPL13A-2xFKBP12::TRP1, RPB1-FRB::KanMX6, [pFR490 (CEN, HIS3, rpb1-CTDS5A-3xFLAG)]* | (Collin et al., 2019) |
| DLY3423 | *Δrrp6,* Rpb1 anchor away + Rpb1-CTD wt-3xFLAG | *as yFR1499, rrp6::URA* | This work |
| DLY3424 | *Δrrp6,* Rpb1 anchor away + Rpb1-CTD S5A-3xFLAG | *as* yFR1503, *rrp6::URA* | This work |
| DLY3417 | *KIN28, ∆rrp6* | *as YDL467, rrp6::KAN* | This work |
| DLY3419 | *kin28 ts16, ∆rrp6* | *as YDL468, rrp6::KAN* | This work |

**Table S13**: plasmids used in this work.

| **Name** | **Description** | Source |
|---|---|---|
| pBS1761 | $Ap^r$, *ori*ColE1; plasmid bearing cassette for N-terminal tagging with PGAL1-TAP | (Finoux and Séraphin, 2006) |
| pDL708 (pU6H3HA) | $Ap^r$, *ori*ColE1; plasmid bearing cassette for C-terminal tagging with *HA* | (Finoux and Séraphin, 2006) |
| pDL772 | $Ap^r$; *ori*ColE1; derivative of pFL38 (URA) bearing yeast *SEN1* | F. Lacroute |
| pDL693 | $Ap^r$; *ori*ColE1; derivative of pFL39 bearing yeast *SEN1* | F. Lacroute |
| pDL703 | $Ap^r$; *ori*ColE1; derivative of pFL39 bearing *sen1ΔNIM* | This work |
| pETM30 | $Kan^r$; oriColE1; vector for overexpression of proteins from the T7 promoter | R. Stefl |
| pDL834 | $Ap^r$; *ori*ColE1; derivative of pFL39 bearing *sen1ΔNter* | This work |
| pDL835 | $Ap^r$; *ori*ColE1; derivative of pFL39 bearing *sen1ΔNterΔNIM* | This work |
| pDL846 | $Kan^r$; oriColE1; derivative of pETM30 carrying the C-terminal domain of Sen1 (aa 1931-2231) under the control of the T7 promoter | This work |
| pDL848 | $Kan^r$; oriColE1; derivative of pETM30 carrying the ΔNIM version of the C-terminal domain of Sen1 (aa 1931-2231) under the control of the T7 promoter | This work |
| pDL856 | $Ap^r$; *ori*ColE1; derivative of pFL39 bearing yeast se*n1-HA* | This work |
| pDL858 | $Ap^r$; *ori*ColE1; derivative of pFL39 bearing *sen1ΔNIM-HA* | This work |
| pDL857 | $Ap^r$; *ori*ColE1; derivative of pFL39 bearing *sen1ΔNter-HA* | This work |
| pDL859 | $Ap^r$; *ori*ColE1; derivative of pFL39 bearing *sen1ΔNterΔNIM-HA* | This work |
| pDL876 | $Ap^r$; *ori*ColE1; derivative of pFL39 bearing the chimeric *nrd1CID* (aa 1-153) - *sen1ΔNter* | This work |
| pDL887 | $Ap^r$; *ori*ColE1; derivative of pFL39 bearing the chimeric *nrd1CID* (aa 1-153) - *sen1ΔNter-HA* | This work |
| pDL965 | $Ap^r$; *ori*ColE1; derivative of pFL39 bearing the chimeric *pcf11CID* (aa 1-137) - *sen1ΔNter* | This work |
| pDL966 | $Ap^r$; *ori*ColE1; derivative of pFL39 bearing yeast se*n1-HTP* | This work |
| pDL967 | $Ap^r$; *ori*ColE1; derivative of pFL39 bearing yeast *sen1ΔNIM -HTP* | This work |
| pDL968 | $Ap^r$; *ori*ColE1; derivative of pFL39 bearing yeast *sen1ΔNter -HTP* | This work |
| pDL991 | $Ap^r$; *ori*ColE1; derivative of pFL39 bearing the chimeric *pcf11CID* (aa 1-137) - *sen1ΔNter-HA* | This work |
| pRS_NC | $Ap^r$; oriColE1; plasmid bearing CID-His6 under the control of the T7 promoter | (Kubicek et al., 2012) |
| pRS_NC_L20D | $Kan^r$; oriColE1; plasmid bearing CID(L20D)-His6 under the control of the T7 promoter | (Vasiljeva et al., 2008) |
| pRS_NC_K21D | $Kan^r$; oriColE1; plasmid bearing CID(K21D)-His6 under the control of the T7 promoter | (Vasiljeva et al., 2008) |
| pRS_NC_S25R | $Ap^r$; oriColE1; plasmid bearing CID(S25R)-His6 under the control of the T7 promoter | (Kubicek et al., 2012) |
| pRS_NC_R28D | $Ap^r$; oriColE1; plasmid bearing CID(R28D)-His6 under the control of the T7 promoter | (Kubicek et al., 2012) |
| pRS_NC_I130A | $Ap^r$; oriColE1; plasmid bearing CID(I130A)-His6 under the control of the T7 promoter | (Tudek et al., 2014) |
| pRS_NC_I130K | $Ap^r$; oriColE1; plasmid bearing CID(I130K)-His6 under the control of the T7 promoter | (Tudek et al., 2014) |
| pRS_NC_I130N | $Ap^r$; oriColE1; plasmid bearing CID(I130N)-His6 under the control of the T7 promoter | (Tudek et al., 2014) |
| pRS_NC_R133A | $Ap^r$; oriColE1; plasmid bearing CID(R133A)-His6 under the control of the T7 promoter | (Tudek et al., 2014) |
| pRS_NC_R133G | $Ap^r$; oriColE1; plasmid bearing CID(R133G)-His6 under the control of the T7 promoter | This work |
| pRS_NC_R133D | $Ap^r$; oriColE1; plasmid bearing CID(R133D)-His6 under the control of the T7 promoter | This work |

**Table S14: List of oligonucleotides used in this work.**

| Name | Sequence (5'-3') | Information/use |
|---|---|---|
| DL1119 | AAGTGACGAAGTTCATGCTA | Forward oligo to generate by PCR a probe to detect snR13 read-through region. |
| DL1367 | GGCCCAACAGTATATTCATATCC | Reverse oligo to generate by PCR a probe to detect snR13 read-through region. |
| DL474 | GCAAAGATCTGTATGAAAGG | Forward oligo to generate by PCR a probe to detect NEL025C. |
| DL480 | ATCTGACCAGGTCAAGCTAC | Reverse oligo to generate by PCR a probe to detect NEL025C. |
| DL2505 | GTGTGTGGACAATCGATTTGC | Forward oligo to generate by PCR a probe to detect snR33 3'precursor and read-through region |
| DL2506 | GCATTGGCTCGATTGTCAAC | Reverse oligo to generate by PCR a probe to detect snR33 3'precursor and read-through region |
| DL1154 | CCTATAACAACAACAACATG | Forward oligo to generate by PCR a probe to detect snR47 RNA. |
| DL1157 | ATAGCCATTAGTAAGTACGC | Reverse oligo to generate by PCR a probe to detect snR47 RNA. |
| DL2627 | ATTCAAAAGCGAACACCGAATTGAC CATGAGGAGACGGTCTGGTTTAT | Reverse oligo used as oligo probe to detect U4 snRNA |
| DL377 | ATGTTCCCAGGTATTGCCGA | Forward oligo to generate by PCR a probe to detect *ACT1* RNA. |
| DL378 | ACACTTGTGGTGAACGATAG | Reverse oligo to generate by PCR a probe to detect *ACT1* RNA. |
